# Identification and characterization of functional modules reflecting transcriptome transition during human neuron maturation

**DOI:** 10.1101/174748

**Authors:** Zhisong He, Qianhui Yu

## Abstract

**Background:** Neuron maturation is a critical process in neurogenesis, during which neurons gain their morphological, electrophysiological and molecular characteristics for their functions as the central components of the nervous system.

**Results:** To better understand the molecular changes during this process, we combined the protein-protein interaction network and public single cell RNA-seq data of mature and immature neurons to identify functional modules relevant to the neuron maturation process in humans. The analysis resulted in 33 discriminable modules which participate in varied functions including energy consumption, synaptic functions and housekeeping functions such as translation and splicing. Based on the identified modules, we trained a neuron maturity index (NMI) model for the quantification of maturation states of single neurons or purified bulk neurons. Applied to multiple single neuron transcriptome data sets of neuron development in humans and mice, the NMI model made estimation of neuron maturity states which were significantly correlated with the neuron maturation trajectories in both species, implying the reproducibility and conservation of the identified transcriptome transition.

**Conclusion:** We identified 33 functional modules whose activities were significantly correlated with single neuron maturity states, which may play important roles in the neuron maturation process.

## Background

As the central organ of the nervous system, the brain is composed of multiple types of neurons and glia ina complex cyto-architecture. By means of synaptic contacts, neurons form local and long-distance networks, which is a key component for brain function. Prior to the establishment of neuronal connections, neurons are generated from neuronal progenitor cells (NPC) located in the areas near the ventricles, and start a long maturation process comprised of a series of sequential and sometimes overlapping steps: neuronal migration, axon elongation, dendrite formation, synaptogenesis and refinement of connections (pruning). This complex developmental process leads immature neurons to eventually acquire their mature appearance and full electrical excitability [1-3]. However, while the molecular changes and regulatory mechanisms of NPC proliferation have been described in detail [4, 5], our knowledge of neuron maturation is still relatively sparse. A comprehensive investigation of neuron maturation at the molecular level could largely expand our understanding not only of brain development and function, but also of neurodevelopmental disorders such as autism and schizophrenia. It could also spark the quantitative measurement of neuronal maturity states, which may provide a powerful tool for future studies.

Here, we adapted an insulated-heat-diffusion-based network smoothing procedure with a topological overlap matrix-based module identification method to analyze differences between immature and mature neurons on the transcriptome level, based on published single-cell RNA sequencing data of adult and fetal human brain tissues and the protein-protein interaction network annotated in the Reactome database. With the identified functional modules discriminating neurons in different maturity states, we developed machine-learning-based neuron maturity indices (or NMIs), which aim to quantify the level of neuron maturity. By applying the NMI models to multiple human and mouse single-cell or purified bulk RNA-seq data from neurons at different developmental stages and conditions, we verified the identified transcriptome transition during neuron maturation in human neuron *in vitro* models, as well as its high conservation in mouse neurons. The constructed NMI models thus show their potential in describing and comparing a variety of neuron maturity states.

## Results

### Detection of protein-protein interaction modules relevant to human neuron maturation

To comprehensively investigate changes of functional modules during the process of neuron maturation in humans, we adapted the module detection algorithm based on the topological overlap matrix (TOM) [6], from the widely used gene co-expression network analysis pipeline WGCNA [7], to the protein-protein interaction network annotated by Reactome [8, 9]. To include gene differential expression information, each edge in the network was weighted by the difference of expression level changes between linked genes, which were smoothed with the insulated heat diffusion procedure to reduce influence of noise (see Materials and Methods). Gene expression level changes during the neuron maturation process in humans were estimated based on the published single cell RNA-seq (scRNA-seq) data of fetal and adult human brains [10].

The analysis resulted in 109 functional modules with sizes ranging from 21 to 203 genes, with a median size of 38 genes (Fig. 1). As shown by the calculated adjusted random index (ARI) [11, 12], choice of insulating parameter did influence the identified modules, but the modular composition remained generally robust (Supplementary Fig. 1). A two-sided Wilcoxon signed rank test was applied to each module in order to identify functional modules with significant expression level changes with concordant direction. Thirty-three functional modules with significant directional changes, which were referred to discriminable modules, were identified (Benjamini-Hochberg (BH) corrected *P*<0.05, Supplementary Table 1). Among them, 17 modules accounting for 964 genes in the network showed higher activity in mature neurons (referred as mature-high modules). On the other hand, the remaining 16 modules accounting for 1125 genes showed higher activity in immature neurons (referred as immature-high modules). Gene Ontology (GO) enrichment analysis by *topGO* [13] and *GOSemSim* [14] indicated that genes encoding for membrane proteins which participate in cell communication, signaling and oxidation-reduction processes for energy generation were strongly enriched in mature-high modules (Fig. 1, Supplementary Table 2). On the other hand, genes encoding for nuclear proteins related to transcription and post-transcriptional processing including splicing and translation were enriched in immature-high modules (Fig. 1, Supplementary Table 2).

**Figure 1.**
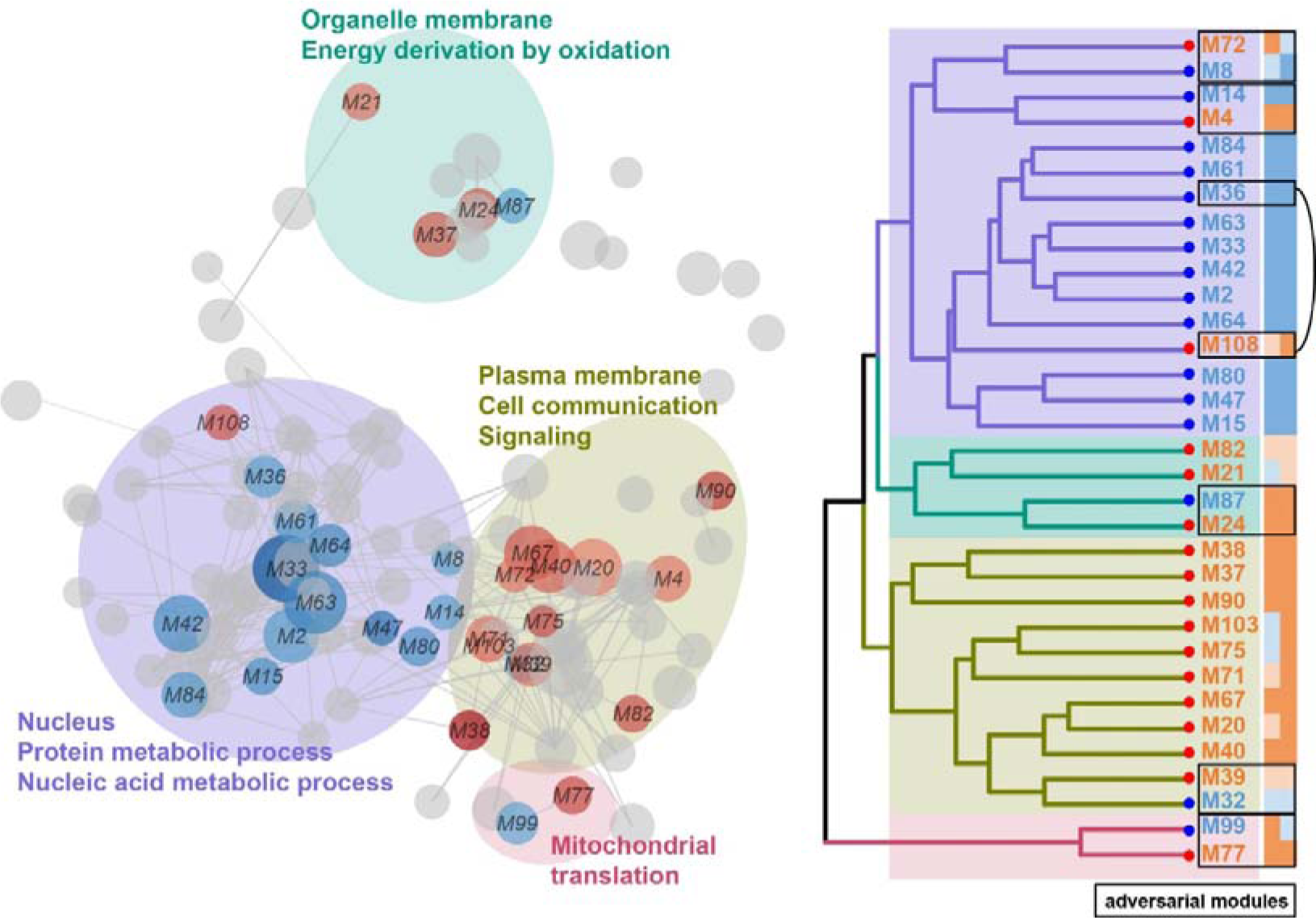
Neuron maturation relevant functional modules in protein-protein interaction (PPI) network. The left panel shows the network of PPI functional modules. Nodes represent distinct modules and are scaled to reflect the number of proteins in each. Colors of nodes represent directions of expression changes during neuron maturation: red – higher in mature neurons, blue – higher in immature neurons, and grey – no significant tendency. Nodes are connected if proteins within the respective modules interact with significantly high frequency. Ellipses mark the functional clusters of modules identified based on hierarchical clustering of Gene Ontology (GO) similarity among discriminable modules using GOSemSim, as shown in the right panel. There, labels are colored to represent directions of expression changes of the respective modules during neuron maturation. Colors of the two columns next to labels show gene expression changes of the respective modules in two brain RNA-seq data (left: He Z, et al. 2014; right: BrainSpan) during prenatal and early postnatal development: red – increase during development, blue – decrease during development. Color darkness indicates whether the change is significant according to Wilcoxon rank sum test. Boxes mark adversarial module pairs.

Although lacking additional data for *in vivo* transcriptome of human neurons across the whole neuron maturation process, it has been reported that neuron maturation explains the majority of brain transcriptome changes during prenatal and new-born postnatal development [15]. Therefore, we took the advantage of fetal and early postnatal brain RNA-seq dataset in BrainSpan and another age series RNA-seq data [16], to compare the brain transcriptome before and after postnatal day 100. Remarkably, 28 out of the 33 discriminable modules showed significant concordant expression level changes (one-sided Wilcoxon signed rank test to fold changes (FC), BH-corrected *P*<0.1) in at least one dataset, while 20 of them showed significant concordance in both datasets (Fig. 1). In addition, although not significant, another two modules showed consistent direction of changes in both datasets. These results suggest that the discriminable modules represent the reproducible functional modules discriminating mature and immature neurons.

### Adversarial functional module pairs

Interestingly, a further comparison with PPI functional modules, which were detected without integrating with expression level differences, identified six adversarial functional module (AFM) pairs. The two modules in one AFM pair were corresponding to the same module when the differential expression information was not integrated (Fig. 1). Three out of six AFM pairs (M4-M14, M36-M108, M8-M72) showed significant expression changes with consistent directions in at least one bulk brain RNA-seq dataset (one-sided Wilcoxon signed rank test, BH corrected *P*<0.01). In addition, consistent discordance in all pairs were observed in both bulk brain datasets (one-sided Wilcoxon rank sum test, *P*<0.01). Further functional analysis revealed highly consistent, connected but varied GO term and biological pathway enrichment in each pair of adversarial modules (*topGO* with the parentchild algorithm for GO terms, one-sided Fisher’s exact test for pathways; BH-corrected *P*<0.05, Supplementary Table 2). This analysis indicated that highly connected biological pathways may play distinct roles during neuron maturation in humans. They may reflect decoupling of components in the same pathway during the neuron maturation process.

A good example is the AFM pair M4-M14 (Fig. 2). Genes in both modules participate in signaling by Rho GTPases, and more specifically, by activating the Rhotekin and Rhophilins pathway according to the Reactome annotation. Interestingly, this pathway splits into two parts: RHOB/C and RTKN in mature-high M4, and RHOA, RHPN1/2 and TAX1BP3 in immature-high M14. This partition implies that, although Rhotekin and Rhophilins both participate in Rho GTPases signaling, they interact with different members of the Rho protein family and play different roles in the process of neuron maturation. Rhophilins interact with RhoA and take part in neuron maturation including neuron migration, which is supported by previous studies suggesting interaction between them [17] as well as the role of Rhophilins in cell migration [18]. Rhetekin, on the other hand, while being important in neural differentiation and neurite outgrowth, is also required for neuron survival [19]. This may explain why the expression level of RTKN gene remains high in mature neurons.

**Figure 2.**
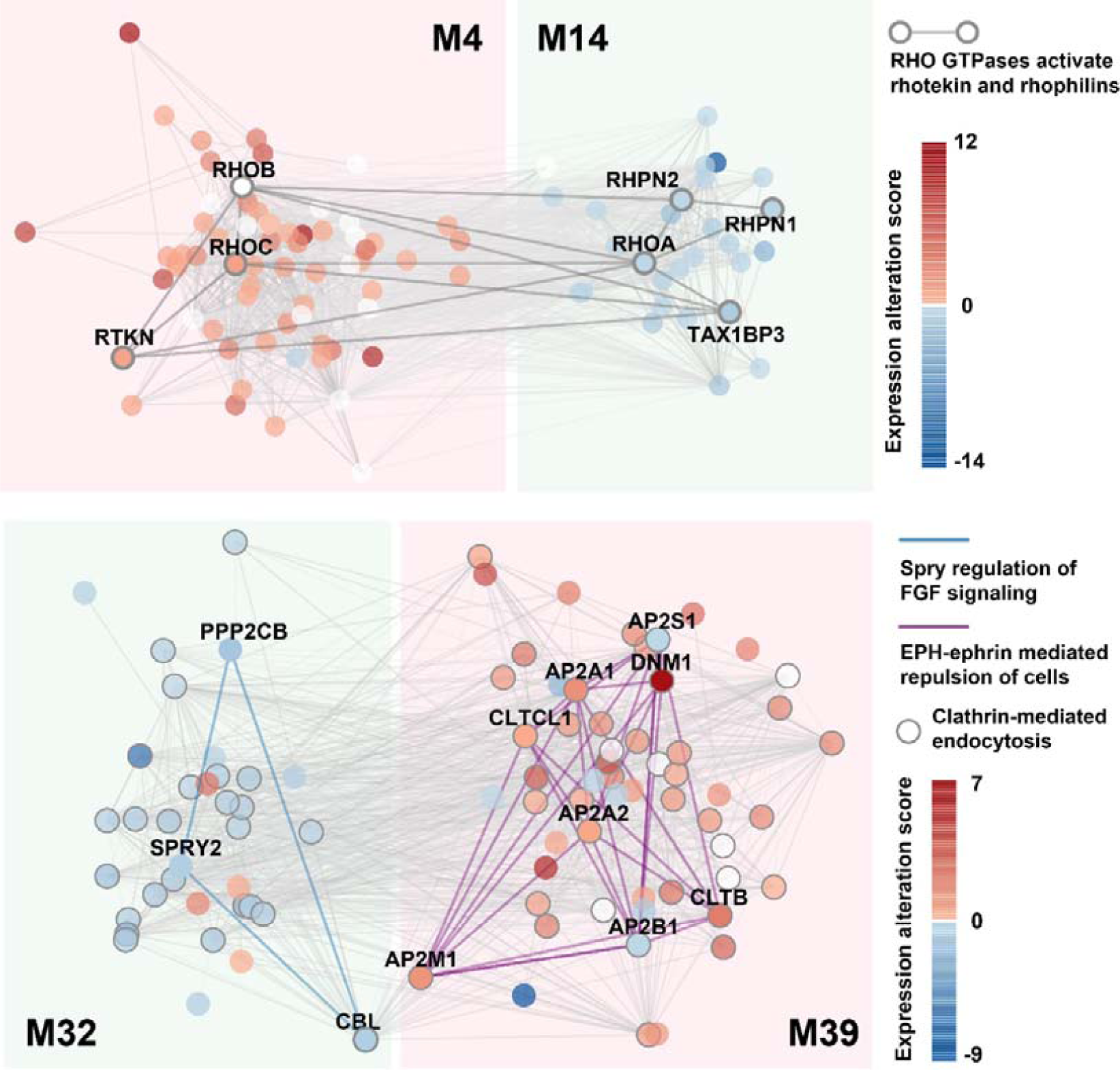
Two examples of adversarial functional module (AFM) pairs. Each circle represents one gene. Edges show annotated PPIs in Reactome among genes in the two functional modules. Colors of circles show expression alteration scores. The upper panel shows the AFM pair M4-M14. Circles with grey border show genes in the network which participate in the pathway “RHO GTPases activates rhotekin and rhophilins”, with PPIs among them shown by the wider grey lines. The lower panel show the AFM pair M32-M39. Circles with grey border show genes which participate in the pathway “Clathrin-mediated endocytosis”. Interactions connecting genes participating in the pathway “Spry regulation of FGF signaling” are shown as blue lines, and interactions connecting genes in the pathway “EPH-ephrin mediated repulsion of cells” are shown as pink lines.

Another AFM pair, M32-M39, represents another scenario. While both modules show significant enrichment of pathways related to endocytosis, genes in the two modules also participate in distinct pathways (Fig. 2). Spry regulation of FGF signaling pathway, which has been reported to be required for cortical development [20], only appears in the immature-high module M32, whereas EPH-ephrin mediated cell repulsion, whose role extends from development to adulthood regulating neuronal plasticity [21], only appears in the mature-high module M39. In summary, the pleiotropy of genes and pathways leads to the separation of the two modules.

### Identified functional modules discriminated different maturity states of neurons from in vitro models

To further estimate how well the neuron-maturation-related transcriptome transitions we identified, especially genes participating in the detected discriminable modules, reflect status of neuron maturation, we establish a machine-learning-based quantitative estimate of neuronal maturity state and tried to apply it to other data sets.

In brief, we constructed a LASSO-regularized logistic regression model based on the standardized expression level of genes involved in each identified module. Each model provided a value ranging between zero and one, namely a modular Neuron Maturity Index (mNMI), with values closer to 1 indicating higher maturity. Ten-fold cross-validation suggested high performances for most of the mNMIs (median AUC=0.87, Supplementary Fig. 2). Applying the models in the test set also resulted in accurate estimations (median AUC=0.84, Supplementary Fig. 2), with those based on discriminable modules performing marginally better (two-sided Wilcoxon rank sum test, *P*=0.11). The mNMIs were further added to two integrated NMIs to represent the overall maturity state, by taking their averages weighted by their performances. This procedure was done by either including all mNMIs (transcriptome NMI or tNMI), or only those based on discriminable modules (discriminable NMI or dNMI). Both general NMIs performed perfectly in distinguishing mature and immature neurons in the test set (AUC=1, Supplementary Fig. 2).

With the NMI models constructed, we applied them to neuron transcriptome data sets of *in vitro* neuron models in order to check whether the identified transcriptome transition could be reproduced and therefore represent the general molecular signature of neuron maturation. In a previous study, Bardy et al. combined patch clamping and scRNA-seq to investigate the relationship between transcriptome and electrophysiology of iPSC-derived neurons [22]. The estimation of NMIs indicated trend of increased neuron maturity accompanying increased action potential, *i.e.* the electrophysiological maturity, especially between the most immature and mature neurons (one-sided Wilcoxon rank sum test, *P*=0.12 for dNMI, *P*=0.02 for tNMI, Fig. 3 and Supplementary Fig. 3).

**Figure 3.**
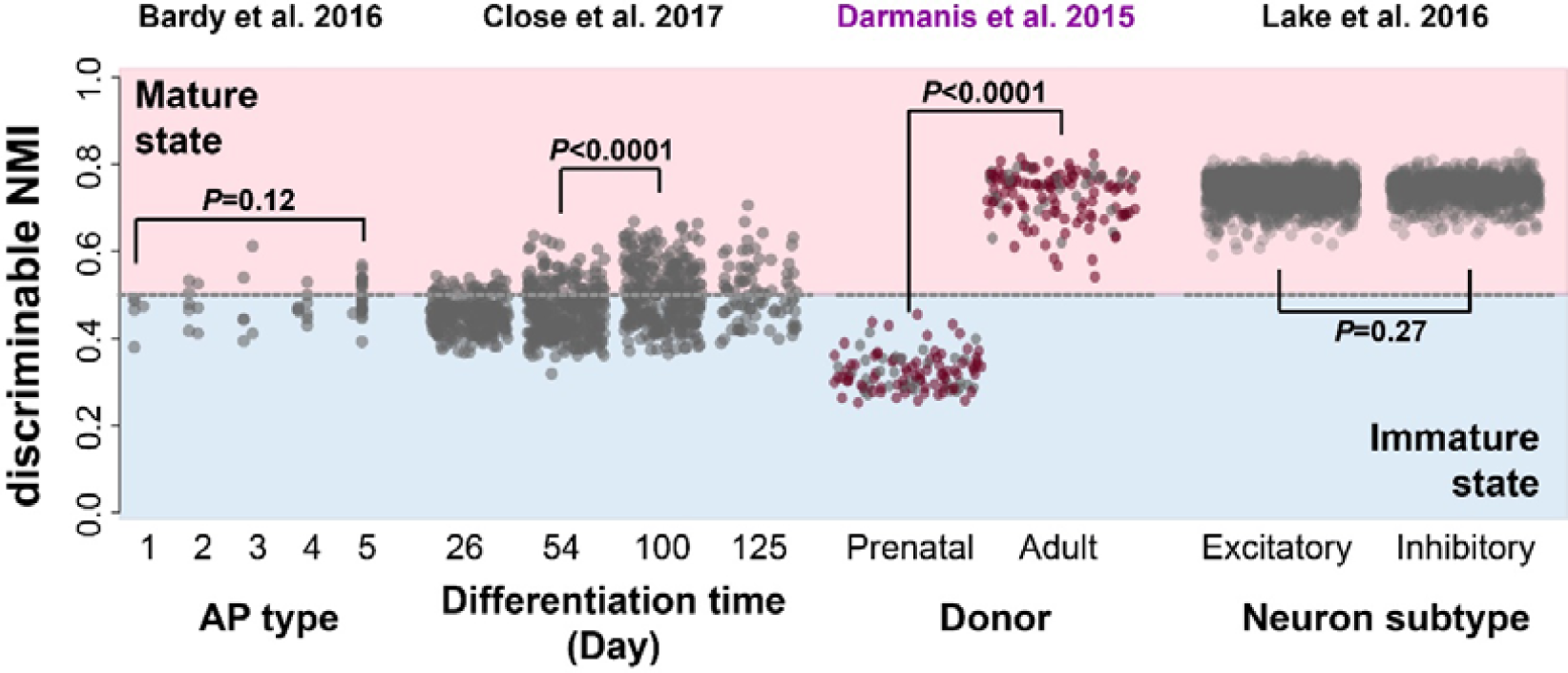
Applications of dNMI in human brain single cell RNA-seq data of neurons to investigate neuron maturity dynamics. The estimated dNMI of each neuron sample is shown, as represented by the y-axis, in four public single cell/nucleus RNA-seq data sets. Each dot represents one cell, with cells involved in the training set colored in brown. The dash line represents NMI=0.5 as the boundary of estimated immature and mature state. For each of the four data sets, cells are grouped based on the respective metadata: Bardy et al. 2016 dataset: action potential (AP) type; Close et al. 2017 dataset: differentiation time; Darmanis et al. 2015: cell donor ages; Lake et al. 2016: neuron subtypes (excitatory and inhibitory neurons). P values of Wilcoxon rank sum test are shown for comparisons of dNMIs between neuron subgroups in each dataset. Purple label on top marks the dataset used to train the NMI model (Darmanis et al. 2015 dataset).

While this dataset was limited by its relatively small number of neurons (*N*=56), Close et al. applied scRNA-seq to interneurons generated by *in vitro* differentiation of human embryonic stem cells (hESCs) to characterize temporal interneuron transcriptome during its maturation, generating another dataset which involved 1733 cells [23]. By estimating NMIs for each DCX^+^ interneuron (*N*=993), we observed the significant increase of integrated NMIs across the time course, especially between 54-day and 100-day (Wilcoxon rank sum test, *P*<0.0001, Fig. 3 and Supplementary Fig. 3). We also noticed that both tNMI and dNMI did not present significant increase between 100-day and 125-day interneurons (Wilcoxon rank sum test, *P*=0.26 for tNMI, *P*=0.58 for dNMI), which is consistent with the weak discrimination between them at the whole transcriptome level proposed by Close et al.

It is worth noting that even at the most electrophysiologically mature state (Bardy et al. dataset) or at the latest time point (Close et al. dataset), a large proportion of interneurons were still in immature state (Fig. 3 and Supplementary Fig. 3). These observations may be due to the technical issue that the NMI model failed to provide prediction of mature neurons, or reflected the failure to complete the neuron maturation process *in vitro*. To answer this question, we examined the human single neuronal nucleus RNA-seq in adult brains [24], resulting in both tNMI and dNMI values significantly larger than 0.5 (Fig. 3 and Supplementary Fig. 3). As expected, no significant difference of both tNMI and dNMI was observed between excitatory and inhibitory neurons (Wilcoxon rank sum test, *P*=0.22 for tNMI, *P*=0.27for dNMI, Fig. 3 and Supplementary Fig. 3). Hierarchical clustering based on Pearson’s correlation coefficient among samples revealed that cell type makes more contributions to sample separation than source of dataset, showing that the estimation is less likely to be biased by batch effect (Supplementary Fig. 4). The above results suggested the potential maturation arrest of the *in vitro* differentiated neurons.

### Transcriptome transition during maturation is conserved in mouse neurons

To check whether the detected transcriptome transition during neuron maturation was conserved in mice, the most widely used animal model for brain development and mental disorders, we applied the constructed human-based NMI model to neuron transcriptome data in mice. Chen et al. extracted maturing interneurons from mouse embryonic medial ganglionic eminence (MGE) and applied scRNA-seq to measure their transcriptome [25]. Estimation of dNMI suggested a boost of maturity state at E17.5, the latest time point across the time course. This result suggested that the human-based NMI models successfully recurred the neuron maturation process in mouse, implying the conserved maturation programs of neuron between humans and mice. Interestingly, the three subtypes of maturing interneurons identified in the study showed significantly different dNMIs (ANOVA, *df*_1_=2, *df*_2_=130, *F*=55.2, *P*<0.0001, Fig. 4A), suggesting that they represented interneurons at distinct stages of maturation.

**Figure 4.**
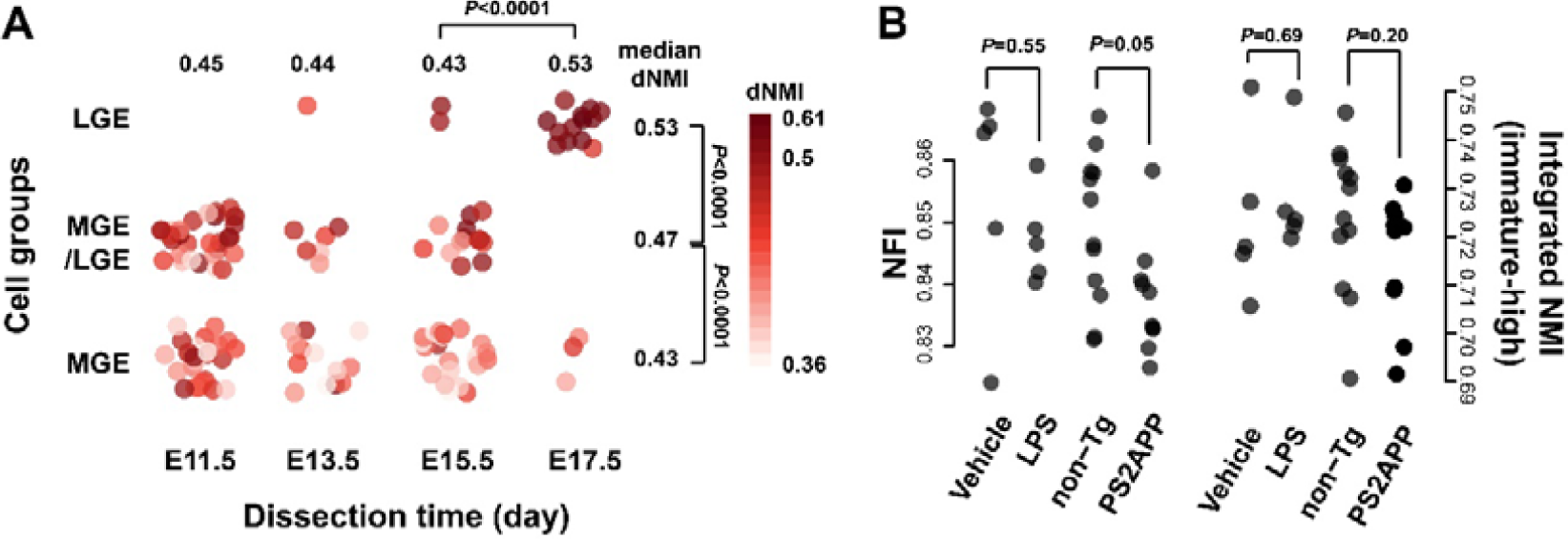
Applications of NMI/NFI in mouse brain neuron RNA-seq data to investigate neuron maturity dynamics. (A) Estimated dNMI of dissected single neurons in mouse medial ganglionic eminence (MGE) based on Chen et al. 2017 dataset. Each dot represents one cell, with color darkness showing maturity states estimated by dNMI. Darker color represents higher level of maturity. Cells are grouped based on the dissection time (x-axis) and cell groups identified by Chen et al. (y-axis). (B) Changes of neuron functionality indicated by neuron functionality index (NFI) in mouse purified neurons responding to neuroinflammation and neurodegeneration, based on Srinivasan et al. 2017 dataset. The left panel shows the estimated NFI, and the right panel shows the integrated NMI of immature-high modules. Each dot represents one purified neuron bulk sample, grouped by the treatment conditions.

### Activities of mature-high modules reflect mature neuron functionality level

Interestingly, applying the dNMI model to the purified neuron transcriptome of PS2APP Alzheimer’s disease mouse model [26] suggested a significantly weaker maturity state than controls (median dNMI_PS2APP_=0.782, median dNMI_control_=0.791, two-sided Wilcoxon rank sum test, *P*=0.003). Further studies on each of the mNMIs indicated that three mNMIs, all of which were based on mature-high modules, significantly decreased in PS2APP neurons (Wilcoxon’s rank sum test, BH corrected *P*<0.1). In addition, among the top-ten of the 27 discriminable modules with reliable mNMIs (AUC>0.8 in cross-validation in training set) and strongest decrease in PS2APP comparing to control neurons, eight were mature-high modules (Fisher’s exact test, odds ratio (OR)=4.25, *P*=0.1). The bias of changes to the mature-high modules was different from observation from the above MGE interneurons data set, as only nine out of 15 (60%) modules with mNMIs significantly different among the three subtypes of maturing interneurons were mature-high modules (Fisher’s exact test, OR=1.83, *P*=0.49).

Considering that the mature-high modules are more likely to be responsible for mature neuronal function maintenance, the biased changes implied that the lower tNMI of PS2APP neurons represented impairment of neuronal function rather than maturation, which has been reported previously [27]. Therefore, we constructed the third integrated index, the neuron functionality index (NFI), which integrated mNMIs from only the mature-high discriminable modules. As expected, the estimated NFIs of PS2APP neurons were significantly lower than those of control neurons (median NFI_PS2APP_=0.836, median NFI_control_=0.850, Wilcoxon rank sum test, *P*=0.05, Fig. 4B). On the other hand, the integrated NMIs of immature-high discriminable modules did not show any significant difference (Wilcoxon rank sum test, *P*=0.58, Fig. 4B). For comparison, no significant difference of either dNMI or NFI was observed between purified neuron transcriptome of a lipopolysaccharide-treated neuroinflammation mouse model and control mouse (Fig. 4B). These results indicated that the activities of mature-high, but not the immature-high, modules may act as signatures of neuron functionality.

## Discussion

In this study, we studied the transcriptome changes during neuron maturation in humans and those functional pathways involved. For this purpose, we developed a new bioinformatics framework, by integrating module identification in the protein-protein interaction (PPI) network and differential expression (DE) analysis. Our strategy revealed 33 functional modules, each of which represents distinct biological pathways, which may be relevant to neuron maturation.

In general, the 17 modules whose genes show significantly higher expression levels in mature neurons, namely mature-high modules, tend to participate in processes relevant to neuronal function and electrophysiology. For instance, there are six discriminable functional modules, all of which are mature-high modules, which show enrichment of synaptic genes and have been reported to be relevant to the electrophysiological maturity of *in vitro* differentiated neurons [22]. Genes in M90, the module enriched for voltage-gated potassium channel complex components, also show higher expression levels in mature neurons. Directly checking those genes in the Bardy et al. dataset suggests higher expression levels in neurons with higher action potential than in neurons with lower action potential in marginal significance (permutation test, *P*=0.052). Furthermore, energy consumption is suggested to grow during neuron maturation, as genes in functional modules related to both respiratory chain (M24) and tricarboxylic acid cycle (M37) show higher expression levels in mature neurons. As previously reported, higher neuronal activity increased mitochondrial oxidative phosphorylation [28]. Therefore, the increasingly active energy generation machinery in mature neurons we observed may be an adaptive strategy of mature neurons to its higher electrophysiological activity.

On the other hand, it is interesting that the 17 immature-high modules whose genes show significantly higher expression levels in immature neurons tend to show enrichment for nuclear functions, which are mainly related to housekeeping processes including RNA and protein metabolism. Indeed, genes in the immature-high modules are significantly overlapped with human housekeeping genes [29], especially when comparing with genes in the mature-high modules (one-sided Fisher’s exact test, odds ratio (OR)=2.1 *P*<0.0001 compared to all genes in the network; OR=3.0, *P*<0.0001 compared to genes in mature-high modules). There are two possible explanations. The decreased activities of housekeeping processes may be an artificial observation due to the increased activities of pathways related to neuronal functions, since the quantification of expression assumes constant amount of transcripts in samples. In such case, genes in the immature-high modules share similar expression level differences which are not relevant to significances of modular expression level differences. However, ANOVA results suggest that genes in different immature-high modules show different amplitude of changes (*F*=6.37, *P*<0.0001). Partial Pearson correlation (PPC) between statistical significances (log-transformed *P*) and modular expression level changes (average expression alteration score) given the module sizes as condition (PPC=0.57, *P*=0.025) suggest dependency between them. Therefore, although this possibility cannot be completely ruled out, there is another scenario, where at least parts of these “housekeeping” processes may play more important roles in the maturation process compared to the final mature stage. This hypothesis is supported by previous studies where mRNA metabolism has proven relevant to some neuronal diseases such as spinal muscular atrophy (SMA) [30], and many regulators of transcription, mRNA translation and protein synthesis have been reported to be related to neurodevelopmental disorders such as autism [31].

To further verify the observed differences between mature and immature neurons, we generated the LASSO logistic regression based neuron maturity index (NMI) model based on the detected gene differential expression, to estimate overall maturity states of neuronal samples. By applying NMI to two public data sets of neurons *in vitro* generated from neuronal progenitor cells (NPC), we find that the constructed NMI model correctly predict neuron maturation statuses. It suggests that the observed transcriptome differences represent general transition during neuron maturation which can be reproduced in neuron models *in vitro*. Meanwhile, we also observed that neurons *in vitro* generated from neuronal progenitor cells (NPC) are likely undergoing maturation arrest, as their estimated maturity states hardly attain complete maturation. In a previous study comparing the transcriptome of *in vitro* neuron models to spatiotemporal human brain transcriptome, *in vitro* neuron models were suggested to be similar to fetal brains [32]. However, the comparison between bulk neural samples with both neurons and proliferative cells can hardly tell whether this similarity is due to the similar NPC:neuron combination, or similar maturity states. Our results suggest that *in vitro* neuronal models are likely to be far from full maturation, which may be due to the lack of environmental stimulation that has been shown to be relevant to neuronal development [33].

Results of applying the NMI model in the mouse medial ganglionic eminence (MGE) single cell RNA-seq data suggests that our observed transcriptome transition happened during neuron maturation is applicable and conserved in mouse. At the same time, it is interesting to see that the three subtypes identified by the study represent neurons with distinct maturity states [25]. In the original study, three neuronal subtypes were identified on a spatial distinction basis: neurons from lateral ganglionic eminence (LGE) expressing LGE markers, neurons from MGE expressing MGE markers, and LGE/MGE neurons expressing both markers. Our study suggests that neurons expressing LGE markers tend to be more mature, and those expressing MGE markers tend to be immature. This observation provides an alternative explanation on a developmental sequential basis, which reconciles with spatial distinction basis explanation, as a previous study has reported that interneurons are generated in MGE and migrate to LGE during their maturation [34]. Together, they provide a more comprehensive description about the origin of interneurons during brain development.

It is worth to mention that our NMI model, although was originally developed to verify the detected transcriptome transition between immature and mature neurons in other data sets, has the potential to corroborate or benchmark transcriptome changes during neuron maturation. Previous studies have developed statistical tools to evaluate maturity state of neural samples, e.g. CoNTExT [32]. Those tools were designed to be used for bulk tissue samples, e.g. dissected brain samples and *in vitro* neural cultures, which consist of multiple cell types including neuronal progenitor cells, immature and mature neurons, as well as non-neuronal glial cells. The NMI model, on the other hand, serves homogeneous neuronal samples, including single neurons and purified neuron populations. In the era of single cell biology, pseudo-time construction analysis, e.g. TSCAN [35], is commonly used to study transcriptome trajectory of cell development, and may be applied also to study neuron maturation [23]. This analysis, however, is limited by lacking benchmark of maturation stages. Although expression of several biomarkers may be helpful in a rough manner, the quantitative description is still missing. The relatively large sample size required to reconstruct a reliable pseudo-time series is also one limitation (although with less significance), as many studies only measured limited number of neurons [22, 25, 36]. In such a scenario, the NMI model can be implemented into, and complement, the existing framework, thus potentially benefitting future research.

## Conclusions

To our knowledge, our study is the first report to comprehensively investigate and characterize molecular functions related to the transcriptome transition happened during neuron maturation in humans. By comparing public single cell RNA-seq data with both immature and mature neurons *in vivo*, we identified 33 functional modules with activities related to neuron maturity states and participating in varied biological processes, including synaptic functions, energy consumption and housekeep processes such as translation and splicing. The detected transcriptome transition was further validated by public human brain transcriptome profiles during development, as well as its high predictive power of neuron maturity states in multiple human neuron *in vitro* models. We also showed that such transition is conserved in mammals, considering its reasonable predictive power of neuron maturity states in mouse neurons.

## Methods

### Identification of neuron maturation relevant functional modules in the human protein-protein interaction (PPI) network

The human protein-protein interaction network was retrieved from the Reactome database (v57) [8, 9], which is comprised of 8,170 proteins and 200,260 undirected interactions. Proteins encoded by genes whose expression was undetectable in brains were excluded, with 5,962 proteins and 125,437 interactions remaining.

Single-cell RNA-seq (scRNA-seq) data of human brains was retrieved from SRA (SRP057196) [10]. The RNA-seq reads were mapped to the human genome hg38 using STAR 2.3.0e [37] with default parameters. The number of reads covering exonic regions of each protein-coding gene annotated in GENCODE v21 was counted and normalized using DESeq2 [38]. FPKM was calculated for each gene in each sample. Average FPKM of each gene was calculated for mature and immature neurons, as the mean FPKM across all cells classified as “neurons” and “fetal quiescent”, respectively. Expression level difference between mature and immature neurons of each gene was represented by expression alteration score *s*:

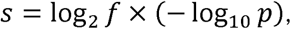

where *f* is the fold change between average FPKM of mature and immature neurons, and *p* is the P-value of ANOVA with neuron maturity state as the independent variable.

A heat-diffusion-based network smoothing procedure, as described and implemented in HotNet2 [39], was then applied to the obtained PPI network where the above expression alteration scores were assigned to corresponding nodes. In brief, a diffusion matrix, which describes the amount of heat diffused between each node pair in the network during the insulated heat diffusion process when the system reaches equilibrium, was defined as:

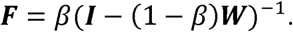

Here, *β* is the insulating parameter (set to 0.55 in this study), and ***W*** is the normalized adjacency matrix. The smoothed expression alteration score of nodes in the network was then calculated as:

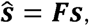

where *s* is the vector of expression alteration scores of all nodes in the network. Weights were assigned to the edges which represent the annotated protein-protein interactions:

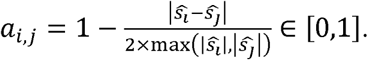

A topological overlap matrix (TOM) based module identification procedure [6], as implemented in WGCNA [7], was then applied to resulted weighted PPI network. In brief, TOM was defined as a *N*×*N*square matrix with *N* as the number of nodes in the network:

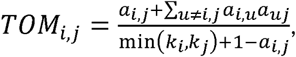

where *a_i,j_* is the weight of edge between node *i* and node *j, k_i_* is the degree of node *i*. Hierarchical clustering with average linkage method was applied using TOM as the distance matrix, followed by the dynamic tree cutting procedure implemented in the R package DynamicTreeCut [40], requiring minimal module size as 20. For each identified module, a Wilcoxon signed rank test was applied to the expression alteration scores of proteins in the module. Modules with Benjamini-Hochberg (BH) corrected *P*<0.05 were defined as discriminable functional modules. Discriminable functional modules with positive median expression alteration scores were defined as mature-high modules, while the remaining ones were defined as immature-high modules.

The pipeline to identify functional modules has been implemented as an R package and can be downloaded at https://github.com/maplesword/TOMRwModule.

### Characterization of functional modules

A Gene Ontology (GO) enrichment analysis was performed for each identified discriminable functional module using the parentChild algorithm [41] implemented in topGO [13], with all genes encoding for proteins involved in the PPI network as background. Pairwise functional similarities of discriminable modules were calculated using GOSemSim [14], by averaging similarities of the three GO categories: cellular component (CC), biological process (BP), and molecular function (MF). Hierarchical clustering with complete linkage was applied to the distances among discriminable modules defined as one minus the calculated similarity.

Functional pathway annotation was performed for each identified discriminable functional module based on the pathway gene set annotation in Reactome using a one-sided Fisher’s exact test to compare with all genes encoding for proteins in the PPI network. Pathways with BH corrected *P*<0.05 were selected.

### Generation of Neuron Maturity Index (NMI)

The NMI models were constructed aiming at the discrimination of mature and immature neurons. To objectively build and test the models, the mature and immature neuron scRNA-seq data mentioned above were randomly separated into two groups. The training set included 99 mature neurons and 82 immature neurons. The test set included 32 mature neurons and 28 immature neurons.

Based on the training set, LASSO logistic regression as implemented in glmnet [42] was applied to each identified functional module, with standardized expression levels of genes in the module in each sample set as independent variables and neuron maturity state as the dependent variable. Expression level standardization was performed for each gene separated as following:

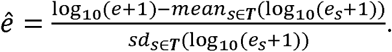

Here, *mean_s∈T_*(log_10_(*e_s_*+1)) and *sd_s∈T_*(log_10_(*e_s_*+1)) represent the mean and standard deviation of the log10-transformed expression levels (in FPKM) across all samples in the training set. The LASSO regularization parameter λ was then determined using ten-fold cross-validation to maximize area under curve (AUC) of receiver operating characteristic (ROC) of the model. For each sample given the expression levels in FPKM, the resulted LASSO logistic regression model of each module predicted the probability of the sample being mature neuron in relative to immature neuron; therefore, it was defined as the modular neuron maturity index (mNMI) of the functional module. Those mNMI models were then applied to the test set for performance evaluation, as well as other neuron scRNA-seq data or purified neuron bulk RNA-seq data for further investigations.

To integrate multiple mNMIs of different functional modules, a weighted mean of multiple mNMIs was calculated for module set ***S***:

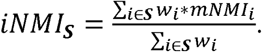

Here, the weight of mNML_*i*_ (*w_i_*) was defined as AUC_i_-0.5, where AUC*_i_* is the AUC of ROC of mNMI during the ten-fold cross-validation in the training dataset. When *S*= *{all functional modules*, the corresponding iNMI_*S*_ was defined as transcriptome NMI (tNMI). Discriminable NMI (dNMI), on the other hand, was defined as the iNMI_*S*_ when ***S***= *{all discriminable modules}*. Lastly, neuron functionality index (NFI) was defined as the iNMI_*S*_ when ***S***=*{all mature-high modules}*.

The NMI models have been implemented as an R package (neuMatIdx) and can be downloaded at https://github.com/maplesword/neuMatIdx.

## Acknowledgements

We thank Prof. Dr. P. Khaitovich, Dr. P. Guijarro and Mr. L. Zhang for the valuable discussions. We thank Prof. Dr. G. Banes and Dr. P. Guijarro for their helpful comments on the manuscript.

## Declarations

### Funding

This work was supported by the National Natural Science Foundation of China (grant 91331203).

### Availability of data and materials

The R package ‘TOMRwModule’ can be downloaded at https://github.com/maplesword/TOMRwModule. The R package ‘neuMatIdx’ can be downloaded at https://github.com/maplesword/neuMatIdx. Complete data necessary to reproduce our results can be accessed at https://maplesword.wixsite.com/home/data.

### Author’s contributions

ZH conceived the project and designed the bioinformatics analysis. ZH and QY executed the bioinformatics analysis and wrote the manuscript.

### Consent for publication

Not applicable

### Ethics approval and consent to participate

Not applicable

### Competing interests

The authors declare that they have no competing interests.

## Abbreviations

AFM: adversarial functional module
ARI: adjusted random index
FC: Fold changes
GO: Gene ontology
NMI: Neuron maturity index
NPC: Neuronal progenitor cells
PPC: Partial Pearson correlation
TOM: Topological overlap matrix

## Supplementary Figures

**Supplementary Figure 1.**
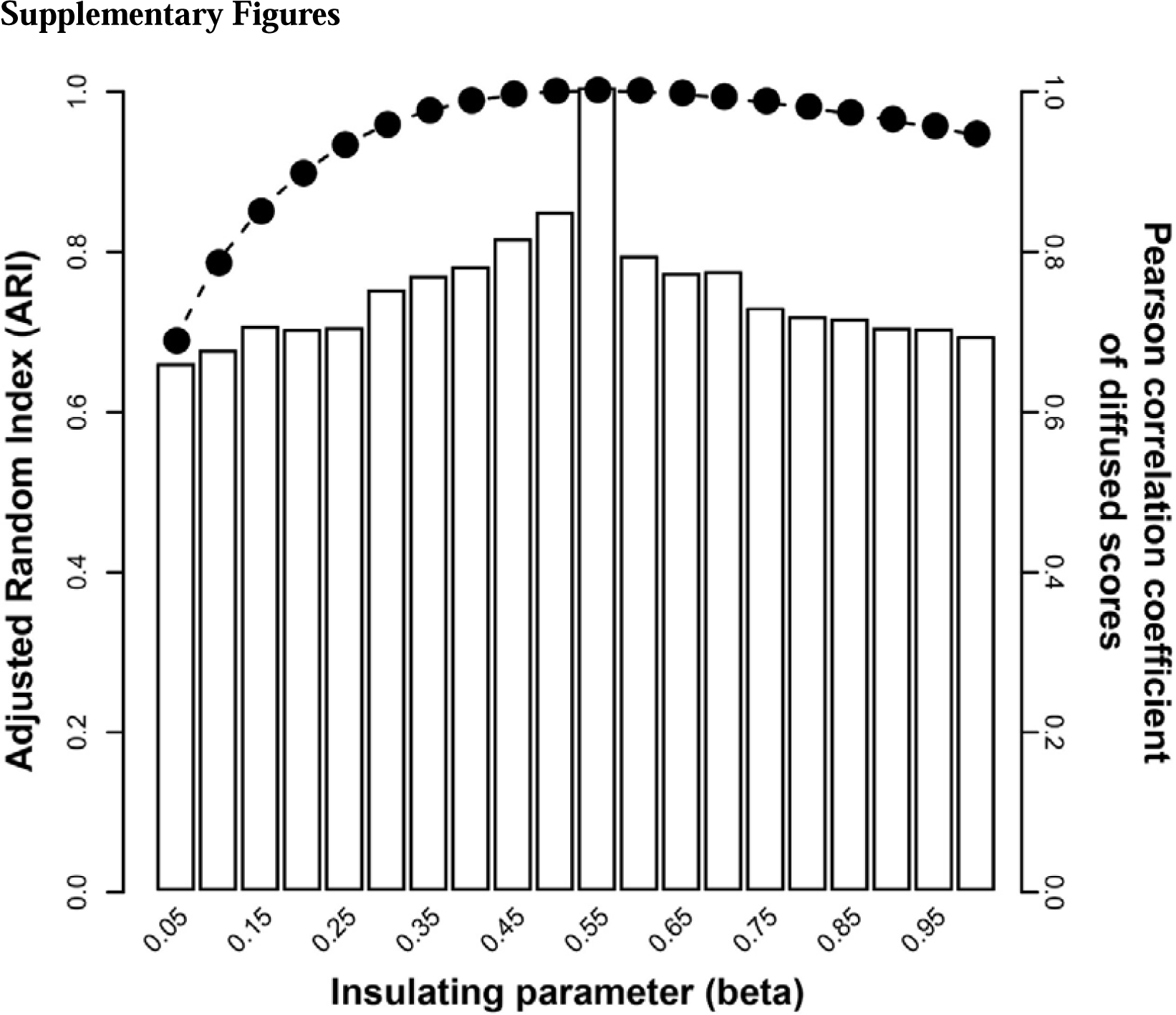
Robustness of module identification to the choice of insulating parameter (β). The x-axis shows 20 different choices of β ranging from 0.05 to 1 with step size of 0.05. Height of each bar shows the adjusted random index (ARI) between modules identified with β=0.55 and those with β set to be the corresponding value as shown by the x-axis. ARI calculates the proportion of agreements between two groupings with adjustment to random performance. Shadow bars show the proportions comparing to random modules. Dots show Pearson correlation coefficient of expression alteration scores after diffusion between choices of β being 0.55 and the value shown by the x-axis.

**Supplementary Figure 2.**
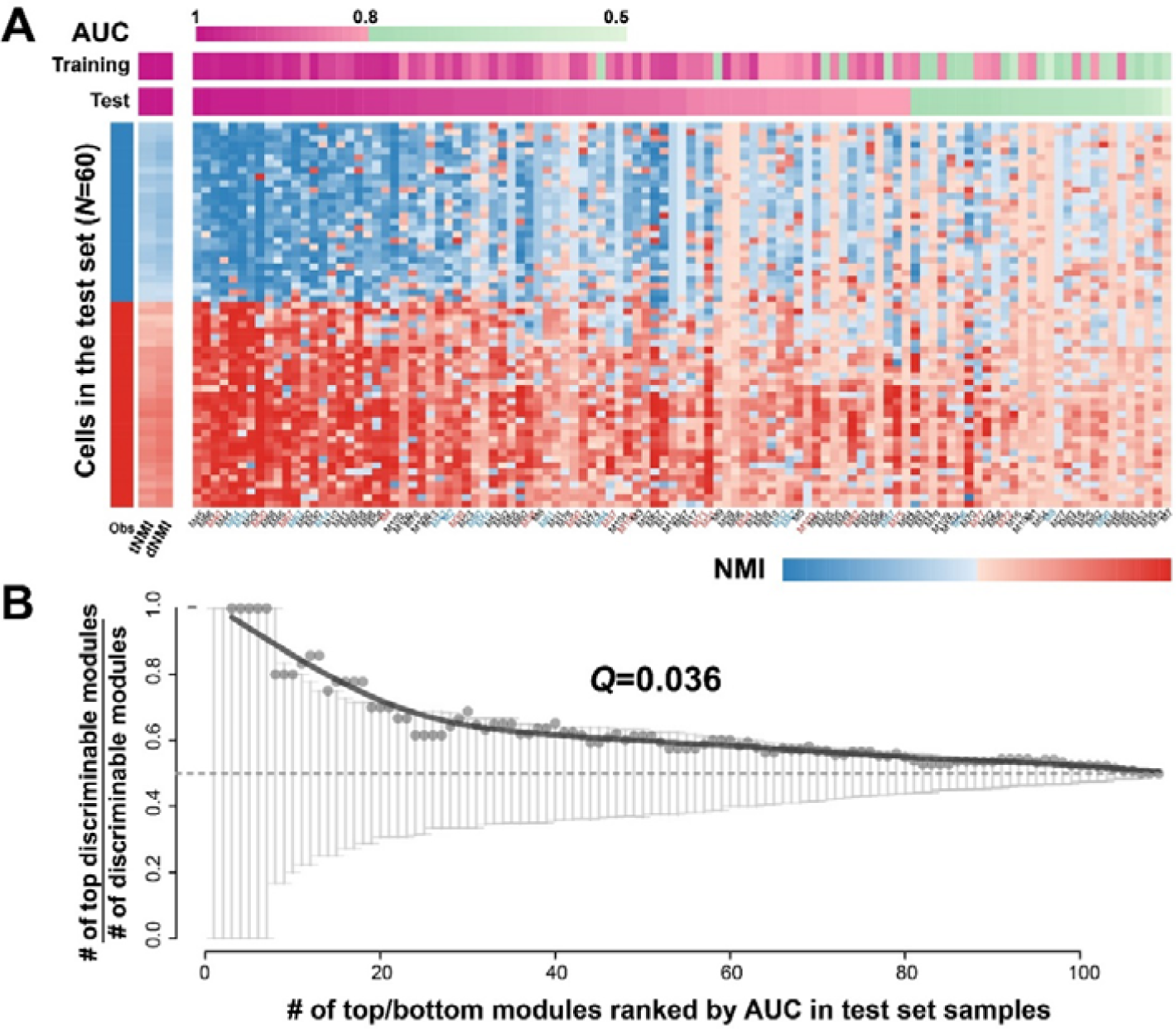
Modular and integrated NMIs of samples in Darmanis et al. dataset. (A) Performance of NMI to estimate neuron maturity state. Bars on top show performance of tNMI, dNMI and each of the mNMIs in prediction of neuron maturity state using Darmanis et al. dataset, as indicated by Area under curve (AUC) of Receiver operating characteristic (ROC). AUC of the training set is calculated based on ten-fold cross-validations. The heatmap shows the estimated NMIs for each neuron in the test set, with each column represent one of tNMI, dNMI and mNMIs of discriminable modules. Module labels are colored based on expression changes of genes in the modules during neuron maturation: red – higher in mature neurons, blue – higher in immature neurons. The real neuron maturity states are shown by the every left column: red – mature neurons, blue – immature neurons. (B) NMIs of discriminable modules perform better than other mNMIs. Y-axis shows ratio between the number of discriminable modules among the top-*N* modules ranked by their mNMI performance in the test set, to the number of discriminable modules among the top-and-bottom-*N* (in total 2*N*) modules. X-axis shows variable *N*. Dots show the observed ratios, with the curve showing the smoothen pattern (natural spline, *df*=5). Grey arrows show the 90% confident intervals based on 1000 permutations of module ranks.

**Supplementary Figure 3.**
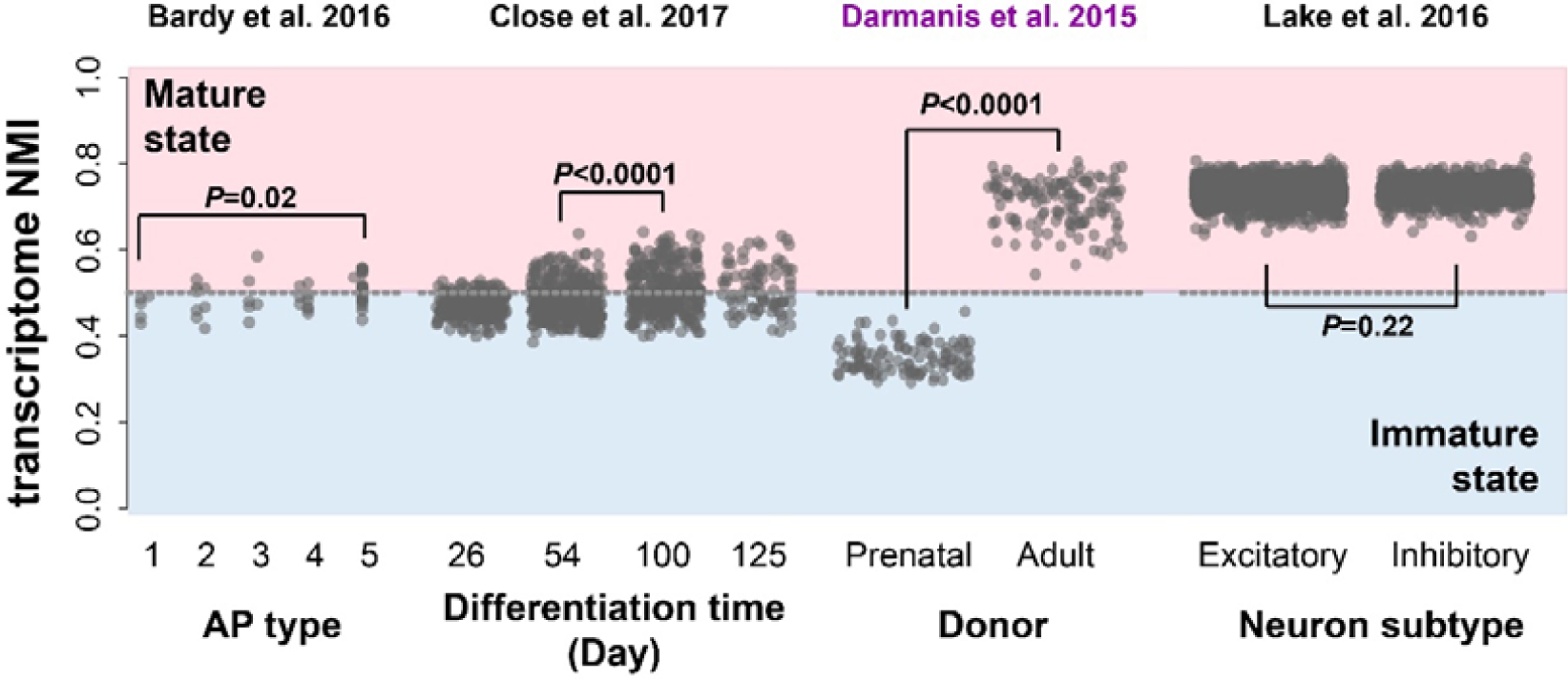
Applications of tNMI in human brain single cell RNA-seq data of neurons to investigate neuron maturity dynamics. It shows the estimated tNMI of each neuron sample, as represented by the y-axis, in four public single cell/nucleus RNA-seq data sets. Each dot represents one cell. The dash line represents NMI=0.5 as the boundary of estimated immature and mature state. For each of the four data sets, cells are grouped based on the respective metadata: Bardy et al. 2016 dataset: action potential (AP) type; Close et al. 2017 dataset: differentiation time; Darmanis et al. 2015: cell donor ages; Lake et al. 2016: neuron subtypes (excitatory and inhibitory neurons). P values of Wilcoxon rank sum test are shown for comparisons of dNMIs between neuron subgroups in each dataset. Purple label on top marks the dataset used to train the NMI model (Darmanis et al. 2015 dataset).

**Supplementary Figure 4.**
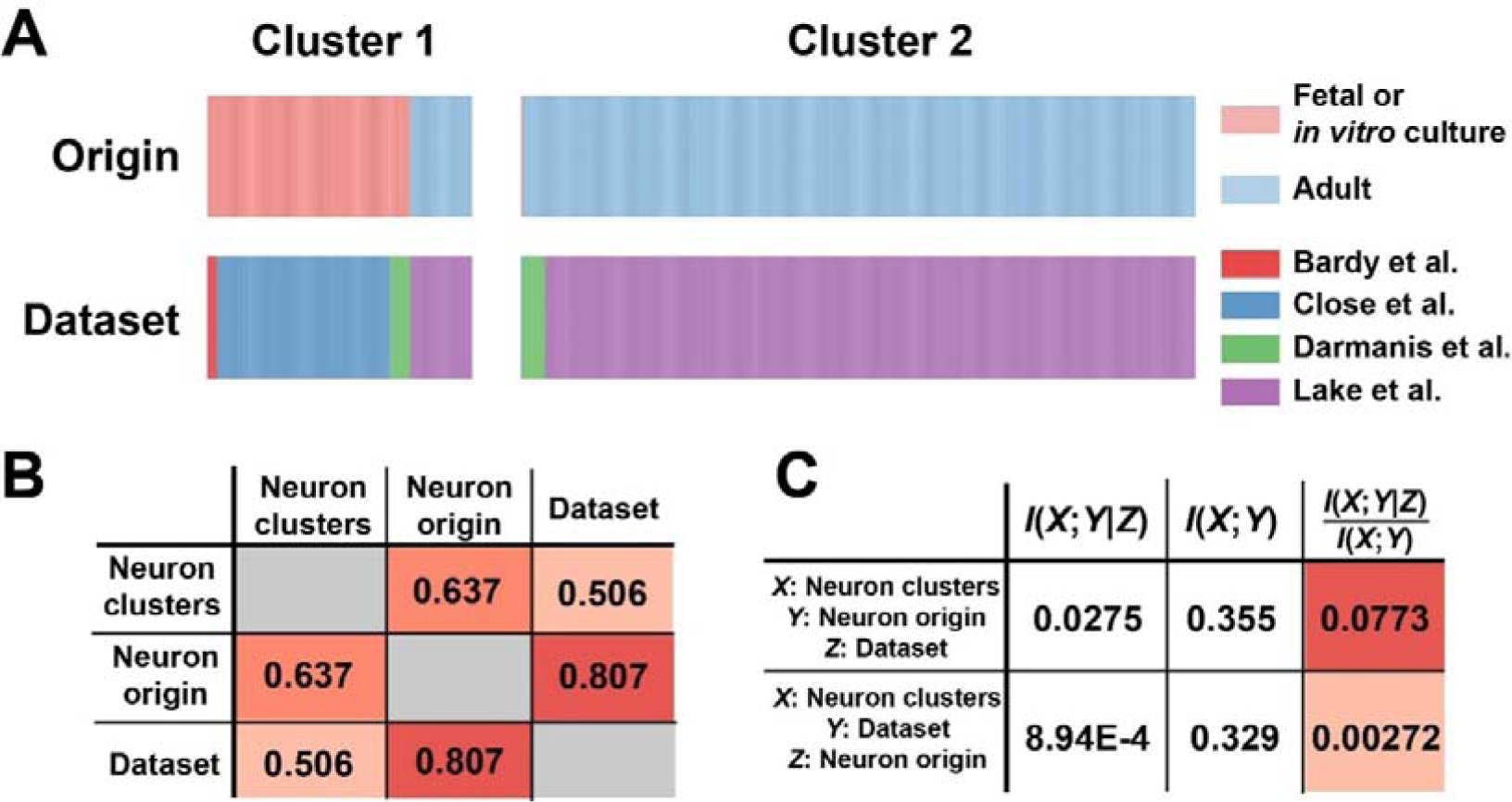
Transcriptome signatures of single neurons are driven by maturity states rather than batch effect across datasets. (A) Origins and datasets of single neurons in the two neuron clusters identified by hierarchical clustering based on standardized expression levels of signature genes for NMI estimations. The two columns show neurons grouped in the two neuron clusters, with the two rows show their origins (red – from fetal brains or *in vitro* cultures) and datasets (red – Bardy et al. dataset; blue – Close et al. dataset; green – Darmanis et al. dataset; purple – Lake et al. dataset). (B) Normalized pairwise mutual information among the neuron clusters, neuron origins and datasets across different single neurons. Darkness of red shows the strength of dependency. (C) Conditional mutual information between neuron clusters and either neuron origins or datasets, under the condition of the other one. The first two columns show the corresponding conditional mutual information and mutual information, with the third column showing the ratio of the first two columns. Darkness of red shows proportions of conditional mutual information among mutual information.

## Supplementary Tables

**Supplementary Table 1**. Characteristics of the identified functional modules

**Supplementary Table 2**. Gene ontology and pathway enrichment of the discriminable functional modules

## References

1. Frank CL, Tsai LH: Alternative functions of core cell cycle regulators in neuronal migration, neuronal maturation, and synaptic plasticity. Neuron 2009, 62(3):312–326.

2. Lalli G: Crucial polarity regulators in axon specification. Essays Biochem 2012, 53:55–68.

3. Rosso SB, Inestrosa NC: WNT signaling in neuronal maturation and synaptogenesis. Front Cell Neurosci 2013, 7:103.

4. Kintner C: Neurogenesis in embryos and in adult neural stem cells. J Neurosci 2002, 22(3):639–643.

5. Urban N, Guillemot F: Neurogenesis in the embryonic and adult brain: same regulators, different roles. Front Cell Neurosci 2014, 8:396.

6. Yip AM, Horvath S: Gene network interconnectedness and the generalized topological overlap measure. BMC Bioinformatics 2007, 8:22.

7. Langfelder P, Horvath S: WGCNA: an R package for weighted correlation network analysis. BMC Bioinformatics 2008, 9:559.

8. Fabregat A, Sidiropoulos K, Garapati P, Gillespie M, Hausmann K, Haw R, Jassal B, Jupe S, Korninger F, McKay S et al: The Reactome pathway Knowledgebase. Nucleic Acids Res 2016, 44(D1):D481–487.

9. Croft D, Mundo AF, Haw R, Milacic M, Weiser J, Wu G, Caudy M, Garapati P, Gillespie M, Kamdar MR et al: The Reactome pathway knowledgebase. Nucleic Acids Res 2014, 42(Database issue):D472–477.

10. Darmanis S, Sloan SA, Zhang Y, Enge M, Caneda C, Shuer LM, Hayden Gephart MG, Barres BA, Quake SR: A survey of human brain transcriptome diversity at the single cell level. Proc Natl AcadSciUSA 2015, 112(23):7285–7290.

11. Hubert L, Arabie P: Comparing partitions. Journal of classificatio 1985, 2:193–218.

12. Lin C, Jain S, Kim H, Bar-Joseph Z: Using neural networks for reducing the dimensions of single-cell RNA-Seq data. Nucleic Acids Res 2017, 45(17):e156.

13. Alexa A, Rahnenfuhrer J: topGO: Enrichment Analysis for Gene Ontology. R package version 2280 2016.

14. Yu G, Li F, Qin Y, Bo X, Wu Y, Wang S: GOSemSim: an R package for measuring semantic similarity among GO terms and gene products. Bioinformatics 2010, 26(7):976–978.

15. Yu Q, He Z: Comprehensive investigation of temporal and autism-associated cell type composition-dependent and independent gene expression changes in human brains. Sei Rep 2017, 7(1):4121.

16. He Z, Bammann H, Han D, Xie G, Khaitovich P: Conserved expression of lincRNA during human and macaque prefrontal cortex development and maturation. RNA 2014, 20(7):1103–1111.

17. Peck JW, Oberst M, Bouker KB, Bowden E, Burbelo PD: The RhoA-binding protein, rhophilin-2, regulates actin cytoskeleton organization. J Biol Chem 2002, 277(46):43924–43932.

18. Nakamura K, Fujita A, Murata T, Watanabe G, Mori C, Fujita J, Watanabe N, Ishizaki T, Yoshida O, Narumiya S: Rhophilin, a small GTPase Rho-binding protein, is abundantly expressed in the mouse testis and localized in the principal piece of the sperm tail. FEBS Lett 1999, 445(1):9–13.

19. Iwai T, Saitoh A, Yamada M, Takahashi K, Hashimoto E, Ukai W, Saito T, Yamada M: Rhotekin modulates differentiation of cultured neural stem cells to neurons. J Neurosci Res 2012, 90(7):1359–1366.

20. Faedo A, Borello U, Rubenstein JL: Repression of Fgf signaling by sprouty1-2 regulates cortical patterning in two distinct regions and times. J Neurosci 2010, 30(11):4015–4023.

21. Kania A, Klein R: Mechanisms of ephrin-Eph signalling in development, physiology and disease. Nat Rev Mol Cell Biol 2016, 17(4):240–256.

22. Bardy C, van den Hurk M, Kakaradov B, Erwin JA, Jaeger BN, Hernandez RV, Eames T, Paucar AA, Gorris M, Marchand C et al: Predicting the functional states of human iPSC-derived neurons with single-cell RNA-seq and electrophysiology. Mol Psychiatry 2016, 21(11):1573–1588.

23. Close JL, Yao Z, Levi BP, Miller JA, Bakken TE, Menon V, Ting JT, Wall A, Krostag AR, Thomsen ER et al: Single-Cell Profiling of an In Vitro Model of Human Interneuron Development Reveals Temporal Dynamics of Cell Type Production and Maturation. Neuron 2017, 93(5):1035–1048 el035.

24. Lake BB, Ai R, Kaeser GE, Salathia NS, Yung YC, Liu R, Wildberg A, Gao D, Fung HL, Chen S et al: Neuronal subtypes and diversity revealed by single-nucleus RNA sequencing of the human brain. Science 2016, 352(6293):1586–1590.

25. Chen YJ, Friedman BA, Ha C, Durinck S, Liu J, Rubenstein JL, Seshagiri S, Modrusan Z: Single-cell RNA sequencing identifies distinct mouse medial ganglionic eminence cell types. Sei Rep 2017, 7:45656.

26. Srinivasan K, Friedman BA, Larson JL, Lauffer BE, Goldstein LD, Appling LL, Borneo J, Poon C, Ho T, Cai F et al: Untangling the brain’s neuroinflammatory and neurodegenerative transcriptional responses. Nat Commun 2016, 7:11295.

27. Sheng M, Sabatini BL, Sudhof TC: Synapses and Alzheimer‘s disease. Cold Spring Harb Perspect Biol 2012, 4(5).

28. Schuchmann S, Buchheim K, Heinemann U, Hosten N, Buttgereit F: Oxygen consumption and mitochondrial membrane potential indicate developmental adaptation in energy metabolism of rat cortical neurons. EurJ Neurosci 2005, 21(10):2721–2732.

29. Eisenberg E, Levanon EY: Human housekeeping genes, revisited. Trends Genet 2013, 29(10):569–574.

30. Linder B, Fischer U, Gehring NH: mRNA metabolism and neuronal disease. FEBS Lett 2015, 589(14): 1598–1606.

31. Kroon T, Sierksma MC, Meredith RM: Investigating mechanisms underlying neurodevelopmental phenotypes of autistic and intellectual disability disorders: a perspective. Front Syst Neurosci 2013, 7:75.

32. Stein JL, de la Torre-Ubieta L, Tian Y, Parikshak NN, Hernandez IA, Marchetto MC, Baker DK, Lu D, Hinman CR, Lowe JK et al: A quantitative framework to evaluate modeling of cortical development by neural stem cells. Neuron 2014, 83(1):69–86.

33. Liu N, He S, Yu X: Early natural stimulation through environmental enrichment accelerates neuronal development in the mouse dentate gyrus. PLoS One 2012, 7(1):e30803.

34. Hansen DV, Lui JH, Flandin P, Yoshikawa K, Rubenstein JL, Alvarez-Buylla A, Kriegstein AR: Non-epithelial stem cells and cortical interneuron production in the human ganglionic eminences. Nat Neurosci 2013, 16(11):1576–1587.

35. Ji Z, Ji H: TSCAN: Pseudo-time reconstruction and evaluation in single-cell RNA-seq analysis. Nucleic Acids Res 2016, 44(13):ell7.

36. Song Y, Botvinnik OB, Lovci MT, Kakaradov B, Liu P, Xu JL, Yeo GW: Single-Cell Alternative Splicing Analysis with Expedition Reveals Splicing Dynamics during Neuron Differentiation. Mol Cell 2017, 67(1):148–161 el45.

37. Dobin A, Davis CA, Schlesinger F, Drenkow J, Zaleski C, Jha S, Batut P, Chaisson M, Gingeras TR: STAR: ultrafast universal RNA-seq aligner. Bioinformatics 2013, 29(1):15–21.

38. Love Ml, Huber W, Anders S: Moderated estimation of fold change and dispersion for RNA-seq data with DESeq2. Genome Biol 2014, 15(12):550.

39. Leiserson MD, Vandin F, Wu HT, Dobson JR, Eldridge JV, Thomas JL, Papoutsaki A, Kim Y, Niu B, McLellan M et al: Pan-cancer network analysis identifies combinations of rare somatic mutations across pathways and protein complexes. Nat Genet 2015, 47(2):106–114.

40. Langfelder P, Zhang B, Horvath S: Defining clusters from a hierarchical cluster tree: the Dynamic Tree Cut package for R. Bioinformatics 2008, 24(5):719–720.

41. Grossmann S, Bauer S, Robinson PN, Vingron M: Improved detection of overrepresentation of Gene-Ontology annotations with parent child analysis. Bioinformatics 2007, 23(22):3024–3031.

42. Friedman J, Hastie T, Tibshirani R: Regularization Paths for Generalized Linear Models via Coordinate Descent. Journal of Statistical Software 2010, 33(1):1–22.

